# Genetic diversity and cross-species transmissibility of bat-associated picornaviruses from Spain

**DOI:** 10.1101/2024.06.20.599728

**Authors:** Marc Carrascosa-Sàez, Jaime Buigues, Adrià Viñals, Iván Andreu-Moreno, Raquel Martínez-Recio, Clàudia Soriano-Tordera, Juan S. Monrós, José M. Cuevas, Rafael Sanjuán

## Abstract

**Background:** Emerging zoonotic diseases arise from cross-species transmission events between wild or domesticated animals and humans, with bats being one of the major reservoirs of zoonotic viruses. Viral metagenomics has led to the discovery of many viruses, but efforts have mainly been focused on some areas of the world and on certain viral families.

**Methods:** We set out to describe full-length genomes of new picorna-like viruses by collecting feces from hundreds of bats captured in different regions of Spain. Viral sequences were obtained by high-throughput Illumina sequencing and analyzed phylogenetically to classify them in the context of known viruses. Linear discriminant analysis (LDA) was performed to infer likely hosts based on genome composition.

**Results:** We found five complete or nearly complete genomes belonging to the family *Picornaviridae*, including a new species of the subfamily *Ensavirinae*. LDA suggested that these were true vertebrate viruses, rather than viruses from the bat diet. Some of these viruses were related to picornaviruses previously found in other bat species from distant geographical regions. We also found a calhevirus genome that most likely belongs to a proposed new family within the order *Picornavirales*, and for which genome composition analysis suggested a plant host.

**Conclusions:** Our findings describe new picorna-like viral species and variants circulating in the Iberian Peninsula, illustrate the wide geographical distribution and interspecies transmissibility of picornaviruses, and suggest new hosts for calheviruses.

## Background

Emerging zoonotic viruses originate from cross-species transmission events involving wild or domesticated animals, particularly mammals, bats being one of the main natural reservoirs of viruses. Several human-infective coronaviruses, filoviruses, paramyxoviruses, lyssaviruses, bunyaviruses, and hantaviruses have a recent bat origin [1–5], and bats might be the source of older cross-species transmissions that led to the evolution of measles, mumps, and hepatitis C [6,7]. Moreover, bats have transmitted viruses of high concern to domestic animals, such as swine acute diarrhea syndrome coronavirus [8]. This prominent role may simply reflect the fact that bats are one of the most abundant and diverse family of mammals, with more than 1200 species described [9,10]. However, it has also been suggested that bats tend to carry more viruses than other mammals due to their highly gregarious and migratory lifestyles, or particularities in their immune system favoring the establishment of persistent infections [11].

The characterization of viruses circulating in wildlife is a daunting task, and although we only know a very small fraction of the virosphere, significant progresses have been made recently [12–14]. Virus discovery has been massively enhanced by metagenomics, which has become the standard approach to describe viral diversity in nature, as well as a powerful surveillance tool [15,16]. The role of bats as likely reservoirs for emerging human coronavirus diseases, particularly COVID-19, has stimulated research focusing on this family of viruses. However, bats are also frequent carriers of other types of viruses that have been less intensively studied, including retroviruses, picornaviruses, herpesviruses and astroviruses [17]. Reflecting this research bias, coronaviruses account for about 50% of the total entries in the Bat-associated virus database [18]. Another important bias in the field is due to differences in research effort around the world, as almost half of the known bat viruses have been reported in China and other parts of Asia.

The Iberian Peninsula is home to more than 30 species of native bats [19], which have been reported to carry lyssaviruses [20], coronaviruses [21], and filoviruses [22], among other types of viruses. Although the Iberian Peninsula and the rest of Europe do not stand out as a global zoonotic hotspot [23], certain risk factors, such as frequent contact between humans and infected animals in disturbed habitats, are likely to occur in this region. In particular, extensive land-use changes and habitat alterations in the Iberian Peninsula may cause disturbances in the distribution of bat colonies and bat-associated viromes, potentially leading to increased spillover risks [24]. In this study, we applied viral metagenomics to bat feces sampled in different regions of Spain and found a relatively high frequency of picornavirus-like sequences. The order *Picornavirales* comprises nine families, including the family *Picornaviridae*, which consists of at least 68 genera containing 158 species, although new groups have been proposed and are currently awaiting approval by the International Committee for the Taxonomy of Viruses (ICTV). Several picornaviruses are pathogenic to humans and other mammals and vertebrates, causing diseases including meningitis, encephalitis, hepatitis, gastroenteritis, myocarditis, paralysis and respiratory infections [25]. Here, we describe five complete or nearly complete genomes belonging to bat-associated picornaviruses from different regions of Spain, as well as a calhevirus belonging to a proposed new family of picorna-like viruses [26].

## Methods

### Sample collection

Fecal samples were collected from 202 individuals belonging to 23 different bat species from May to October 2022 in different Spanish regions (Cantabria, Castellón, Lugo, Murcia, Salamanca, Teruel, and Valencia; **Figure 1**). Bats were captured using nylon mist nets and a harp trap (Austbat), and individuals were briefly placed in cotton bags to recover fresh fecal samples. Then, samples were collected separately for each individual, placed in tubes containing 500 µL of 1X phosphate-buffered saline (PBS), kept cold initially, and then at −20 °C until they were transported to the laboratory and stored at −80 °C.

**Figure 1.**
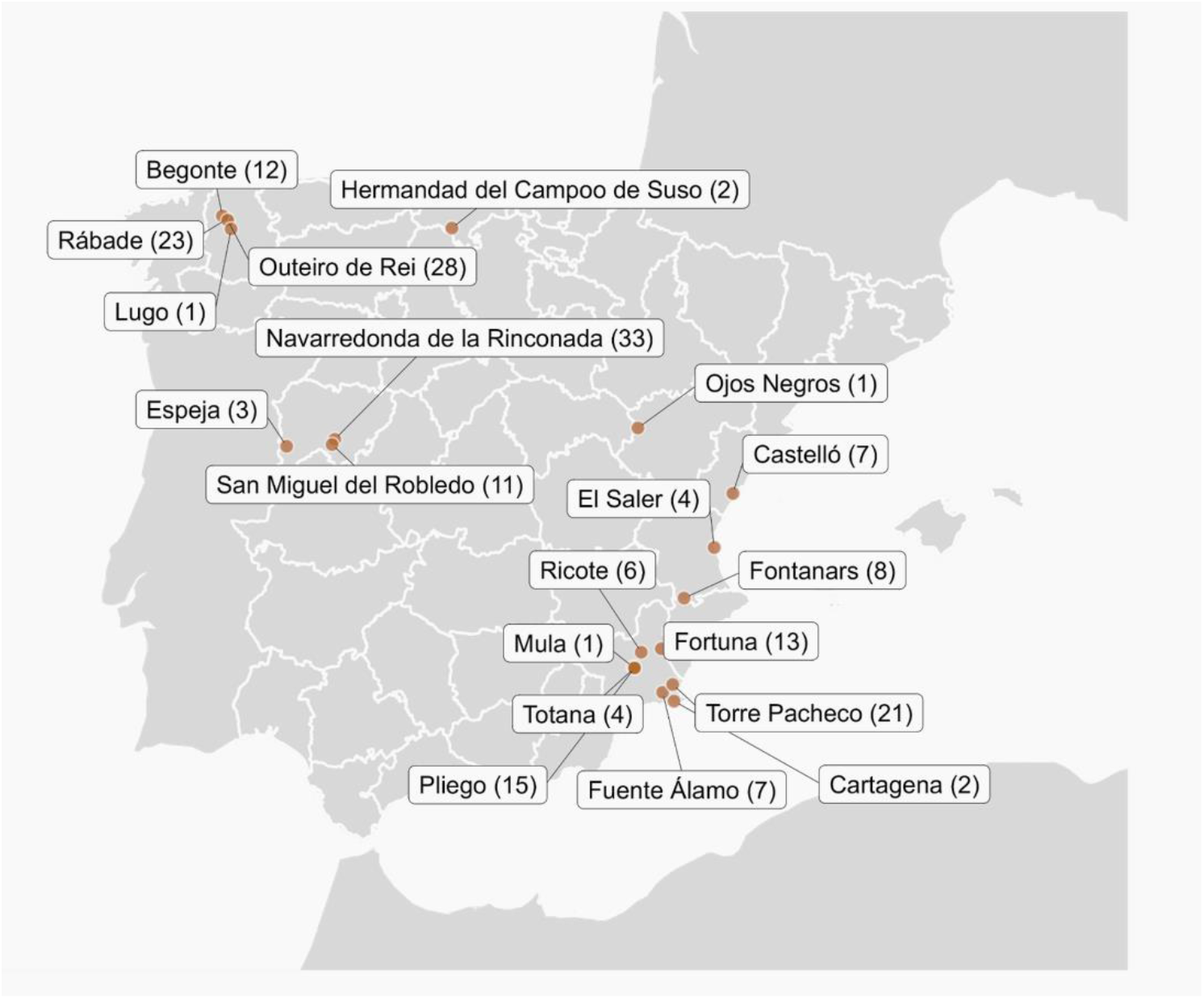
Bat captures for virome analysis. Names of the localities where sampling was carried out and the number of fecal samples collected are indicated.

### Sample processing and RNA extraction

A fraction of the collected samples was combined into a total of 26 pools, each including between one and 15 samples from the same bat species (**Table S1**). Before starting sample processing, each pool was spiked with 10^5^ plaque-forming units (PFU) of vesicular stomatitis virus (VSV) to assess final viral recovery efficiency. Then, each pool was homogenized in a Precellys Evolution tissue homogenizer (Bertin) using 2 mL tubes and 1.4 mm ceramic beads. The homogenization protocol consisted of three cycles of 30 s each at 6500 rpm, with a 10 s pause between cycles. After homogenization, samples were centrifuged twice at 20,000 g for 3 min at 4 °C. Supernatants were then filtered using Minisart cellulose acetate syringe filters with a 1.2 µm pore size (Sartorius) and transferred to ultra-clean 2 mL tubes. For RNA extraction, 250 µL of the filtrate were initially cleaned with Trizol LS reagent (Invitrogen) and then 280 µL from this first step were used for final extraction using the QIAamp Viral RNA minikit (Qiagen). RNA was eluted in a final volume of 40 µL and stored at −80 °C.

### Sequencing and classification of viral sequences

Extracted RNA was used for library preparation using the stranded mRNA preparation kit (Illumina), skipping mRNA enrichment steps, and starting at the fragmentation step. Samples were subjected to paired-end sequencing using a NextSeq 550 device with read length of 150 bp at each end. Raw reads were de-duplicated and quality filtered using fastp v0.23.2 [27]. A quality trimming threshold of 20 was used, and reads below 70 nt in length were removed from the data set using fastp v0.23.2. De novo sequence assembly was performed using SPAdes v3.15.4 [28] with the meta option, and MEGAHIT v1.2.9 [29] with default parameters. Then, contigs assembled with both methods were clustered to remove replicates or small replicates of larger contigs using CD-HIT v4.8.1 [30], and those shorter than 1000 nt were removed. The obtained clustered sequences were taxonomically classified using Kaiju v1.9.0 [31] with the subset of NCBI nr protein database downloaded on June 6, 2023. Clustered sequences were also analyzed using Virsorter2 v2.2.4 [32] to detect viral contigs. Then, viral contigs were analyzed with CheckV v1.0.1 [33] using the CheckV database v1.5 to further assess their quality. Finally, contigs assigned to the *Picornaviridae* family or associated with picornaviruses were selected.

### Genome annotation and phylogenetic analysis

Sequences similar to each contig of interest were searched using DIAMOND v2.0.15.153 [34] with the blastx option and the NCBI nr database downloaded on June 7, 2023. Then, the 100 sequences closest to each contig were extracted and analyzed for association with vertebrate-infecting viruses, while protein domains were annotated using Interproscan v5.63-95.0 [35] using the Pfam database v35.0. Open reading frames (ORFs) were predicted using ORFfinder. Nucleotide-based multiple alignments were performed using MAFFT v7.490 [36] and, where indicated, pairwise identities were calculated using the Sequence Demarcation Toolkit (SDT) v1.2 [37]. Coverage statistics of selected viral contigs were calculated by remapping the trimmed and filtered reads to each contig using Bowtie2 v2.2.5 [38]. In addition, selected viral contigs were compared to NCBI databases using BLAST to obtain identity values and refine annotations. For phylogenetic analysis of P1 and 3D^pol^ regions, nucleotide sequences were aligned with Clustal Omega v1.4.1 [39] and trimmed with ClipKIT [40]. The best substitution model was inferred using the ModelFinder feature in IQTREE [41], which was the GTR+I+G4 model for the P1 and 3D^pol^ *Picornaviridae* family phylogenies, GTR+I+G4+F for the whole polyprotein *Picornaviridae* family phylogeny, and Rtrev+G4 for the *Picornavirales* phylogeny. Phylogenetic tree inference was performed using Mr.Bayes v3.2.7 [42] or IQ-TREE (v2.0.3) [43]. Branch support was estimated with 10,000 ultra-fast bootstrapping replicates and 1,000 bootstrap replicates for the SH-like approximate likelihood ratio test.

### PCR confirmation

The origin of the new viruses detected using metagenomics was determined by individual PCR analysis of the different samples included in each pool. For this purpose, RNA isolation was independently performed for each sample with the QIAamp Viral RNA mini kit (Qiagen) following manufacturer’s instructions. RNA was eluted in a volume of 30 µL, and 4 µL were used to obtain first strand cDNA with Superscript IV reverse transcriptase (Invitrogen) using specific reverse primers for each virus. Then, amplification was performed using NZYTaq II Green Master Mix (NZYTech) and specific primers (**Table S3**). PCR products were visualized by electrophoresis on a 1% agarose gel using Green Safe Premium (NZYTech).

### Nucleotide composition analysis

Nucleotide composition analysis was performed as previously described [26,44]. Briefly, picorna-like genome sequences were downloaded from Virus-Host DB [45] on December, 2023 (www.genome.jp/virushostdb). Viruses defined as infecting more than one host type (i.e. mammals, insects, or plants) were removed and those with segmented genomes were concatenated, resulting in a total of 1065 genomes. Mononucleotide frequencies, dinucleotide frequencies and dinucleotide bias were calculated using the Biostrings package v.2.70.1 in R v4.3.2. These three sets of metrics were used to predict viral host through linear discriminant analysis using the R MASS package. Finally, the predictive power of the model was tested by leave-one-out cross-validation. Accession numbers, reported host, mono- and dinucleotide frequencies, dinucleotide bias, and resulting linear discriminants are reported in **Table S4**.

## Results and Discussion

### Overview

We obtained droppings from 202 animals belonging to 23 different bat species and captured in seven different regions of Spain between May and October 2022 (**Figure 1**; **Table S1**). Fecal material from individuals of the same species was pooled before sample processing. After RNA extraction and library preparation, Illumina sequencing generated between 4.5 and 48 million raw reads per pool (**Table S2**). Quality-filtered reads were de novo assembled and the resulting contigs were analyzed to detect viral sequences. In total, 430 high-quality complete or nearly complete viral genomes were obtained, the vast majority of which corresponded to bacteriophages. Of these, six corresponded to picorna-like sequences of at least 7kb, representing complete or quasi-complete viral genomes. These included five picornaviruses genomes, while the sixth was a new calhevirus. Proposed names for these viruses are provided in **Table 1**. Since the pools analysed contained samples from different regions, RNA extraction and RT-PCR were performed for each sample (**Table S3**) in order to assign the geographical origin of described picorna-like genomes.

**Table 1.** Summary of picorna-like viral genomes found in bat feces from Spain.

| Proposed name | Lugo bat picornavirus | Navarredonda bat picornavirus | Almagra bat picornavirus | Cabezo Gordo bat-associated mischivirus | Fontanars bat-associated kobuvirus | Castelló calhevirus |
| --- | --- | --- | --- | --- | --- | --- |
| Accession | PP654851 | PP654853 | PP654854 | PP654855 | PP654852 | PP654856 |
| Size (kb) | 7.0 | 7.7 | 7.7 | 8.5 | 8.5 | 10.1 |
| Mean coverage | 105.6 | 95.5 | 31.3 | 222.7 | 20036.2 | 692.2 |
| Bat species | *Barbastella barbastellus* | *Myotis myotis* | *Myotis blythii* | *Miniopterus schreibersii* | *Pipistrellus kuhlii* | *Tadarida teniotis* |
| Location | Rábade (Lugo), Begonte (Lugo) | Navarredonda de la Rinconada (Salamanca) | Cueva de la Almagra (Fortuna, Murcia) | Cabeza Gordo (Torre Pachecho, Murcia) | Fontanars (València) | Castelló (Castelló) |
| Proposed virus species | Lugo bat picornavirus | Scotophilus kuhlii picornavirus | Scotophilus kuhlii picornavirus | mischivirus B1 | Wenzhou bat picornavirus 7 | Castelló calhevirus |
| Closest Genbank accession | Bat picornavirus KJ641689.1 | Scotophilus kuhlii picornavirus MZ544240.1 | Scotophilus kuhlii picornavirus MZ544245.1 | Bat picornavirus KP054278.1 | Wenzhou bat picornavirus 7 OQ716046.1 | Jingmen bat picorna-like virus OQ716071.1 |
| Amino acid identity (coverage) | 75.4 (37)% | 78.9 (99)% | 78.9 (99)% | 87.6 (80)% | 74.9 (29)% | 68.7 (28)% |

We then annotated and characterized the genome structure of each picornavirus (**Figure 2**), analysed characteristic sequence motifs (**Table 2**), and performed a Bayesian phylogenetic analysis to define their relationship to those of ICTV-approved viral species, using both the P1 and 3D^pol^ sequences for the *Picornaviridae* members (**Figure 3**) and the RdRp sequence for the calhevirus (**Figure 4**). However, since most viral genomes have not yet been officially classified, we also looked for the most similar sequences in Genbank to ascertain whether our newly described viruses constituted novel species according to ICTV picornavirus species demarcation criteria [46]. Finally, in an attempt to rule out a diet or environmental origin, we used linear discriminant analysis (LDA) to infer the most likely host of each virus based on nucleotide composition and dinucleotide biases (**Figure 5, Table S4**). Using 1065 picorna-like viruses (794 mammalian, 34 fish, 59 avian, 99 plant, and 79 insect viruses), leave-one-out cross-validation indicated that vertebrate, plant, and insect viruses can be confidently distinguished, whereas the separation between mammalian, avian and fish viruses is less clear (**Figure 5A**). This model was then applied to identify the most likely host of the newly described bat-associated viruses. For the sake of clarity, the following sections will detail the above analyses individually for each new virus described.

**Figure 2.**
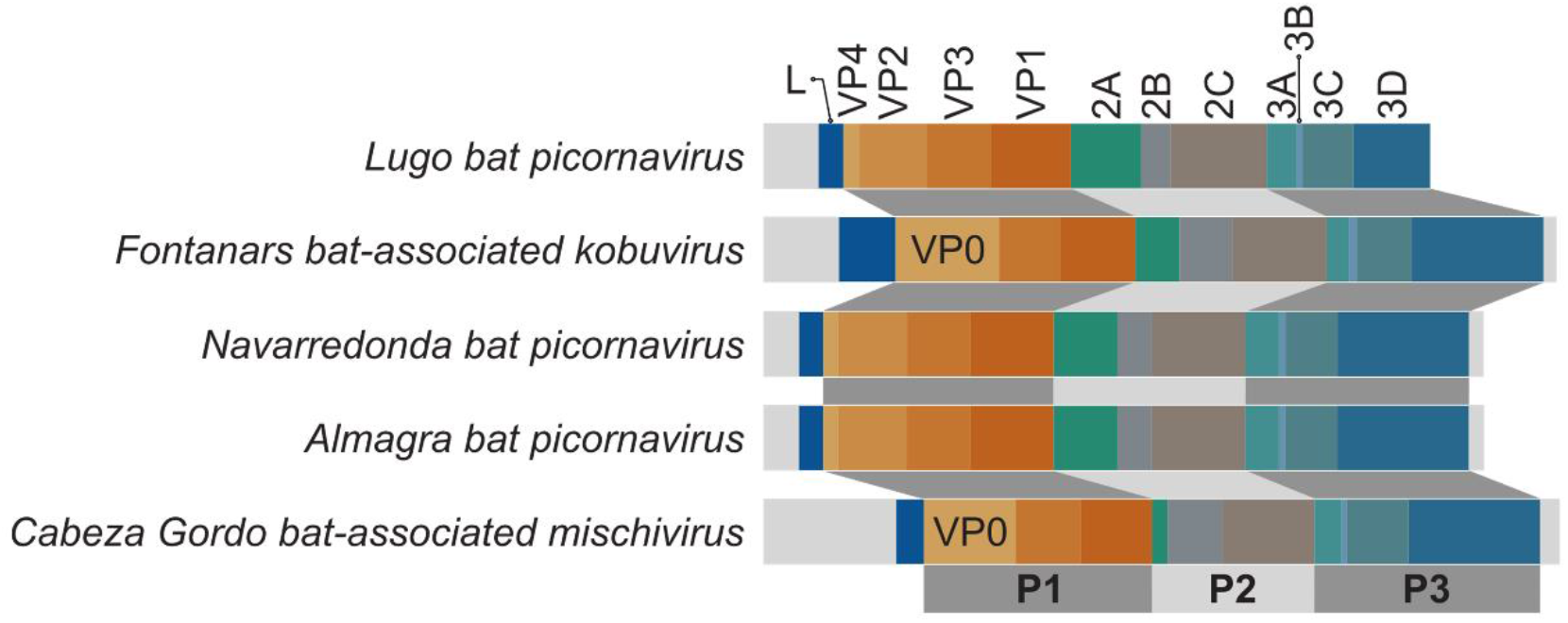
Genome organization of viruses identified in this study belonging to *Picornaviridae* family. Open reading frames for each mature protein (L, VP1-4, 2A-C, 3A-D) are shown as colored regions and mapped to P1, P2, or P3 polyproteins accordingly.

**Figure 3.**
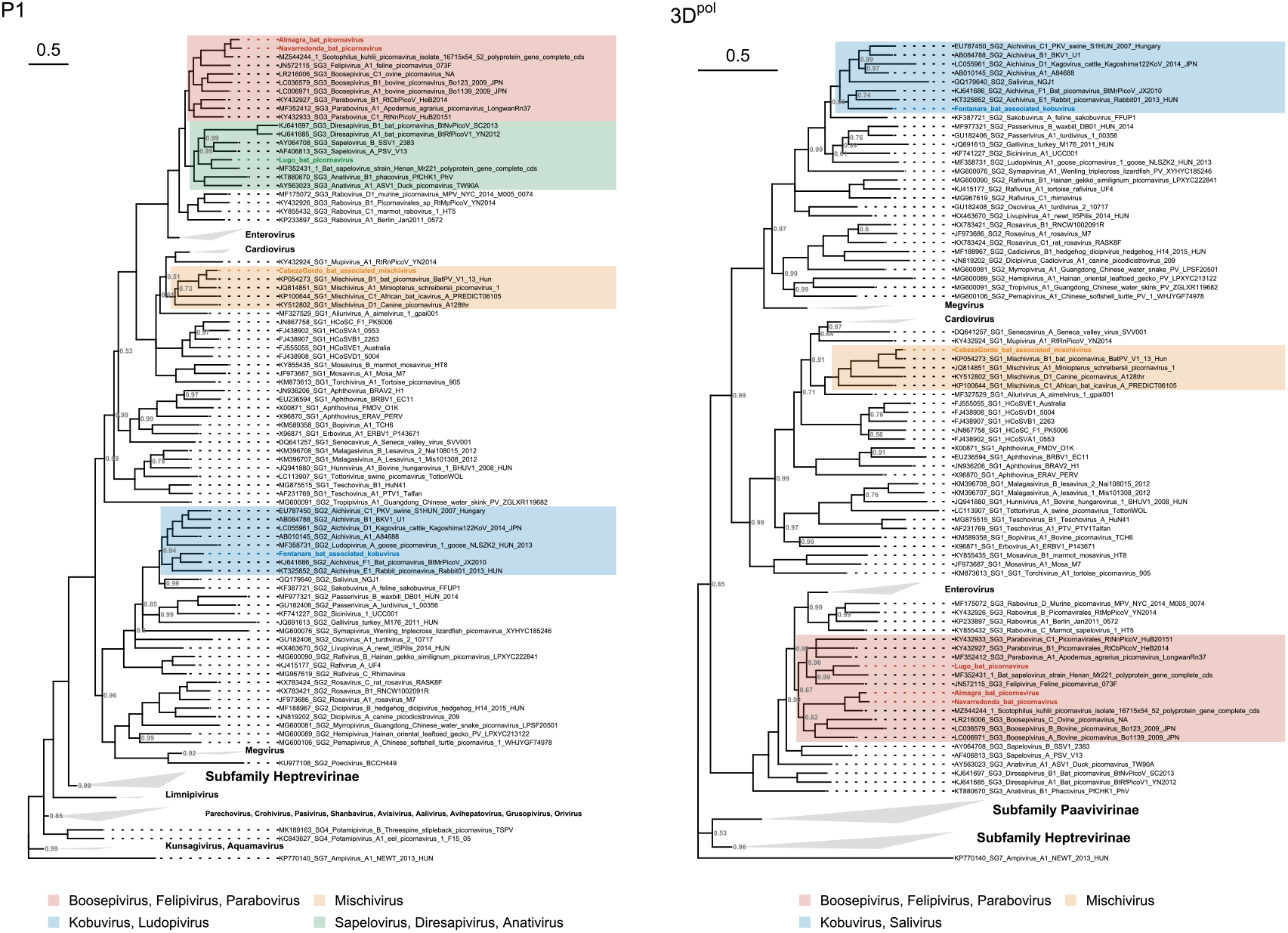
Phylogenetic trees of the *Picornaviridae* family for the P1 and 3D^pol^ regions. The newly described viruses are placed in the context of ICTV-approved viruses. Ampivirus A1 (accession KP770140) was selected as an outgroup for both trees. Phylogenetic inference was performed with the Bayesian MCMC algorithm using a GTR+I+G4 substitution model, and 12 and 8 million generations for P1 and 3D^pol^ phylogenies, respectively. Posterior probabilities for each node are indicated (values of 1 are omitted).

**Figure 4.**
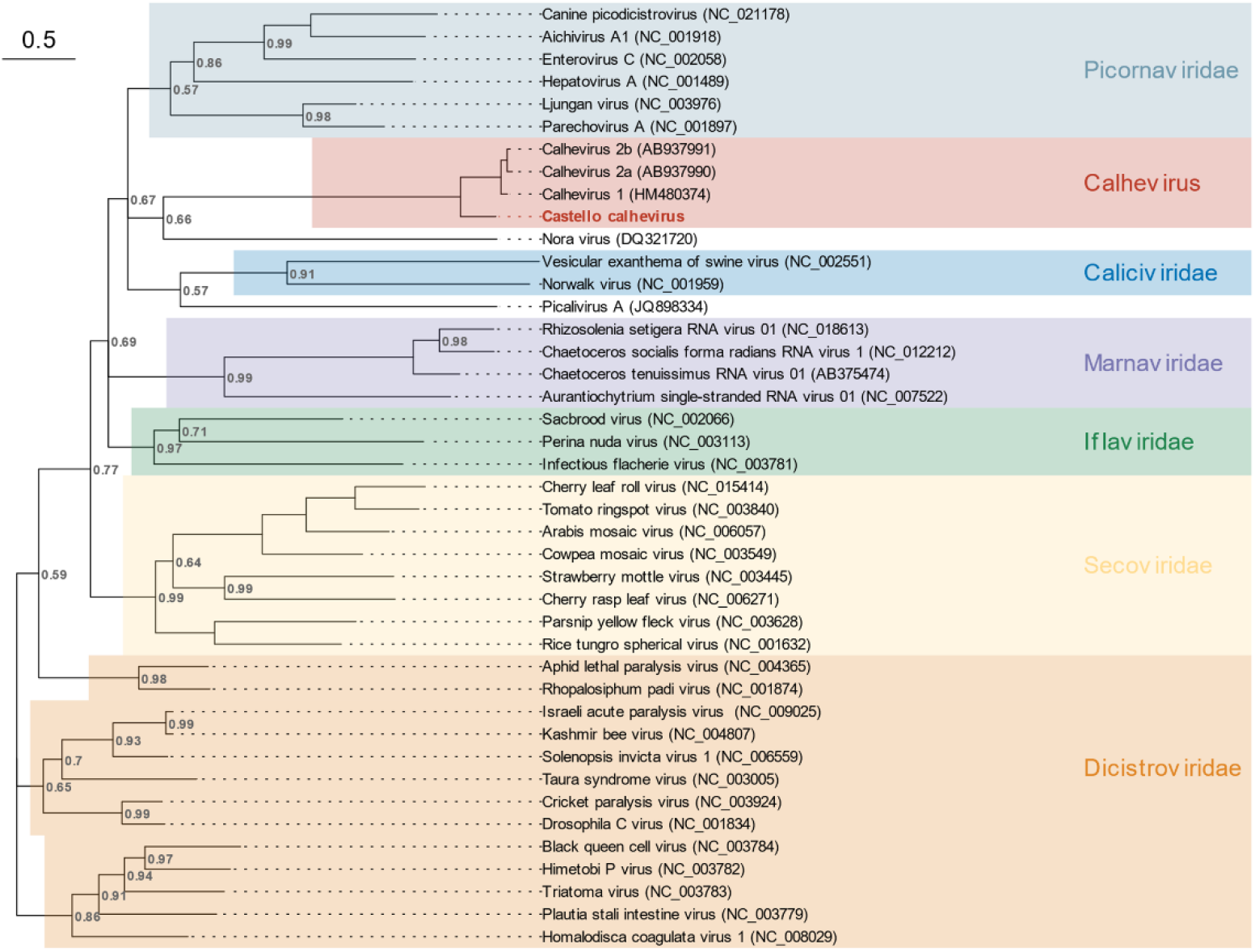
Phylogenetic analysis of the RdRp region for *Caliciviridae*, *Dicistroviridae*, *Iflaviridae*, *Marnaviridae*, *Noraviridae*, *Picornaviridae*, and *Secoviridae* families, previously known calheviruses, and Castelló calhevirus. Phylogenetic inference was performed with the Bayesian MCMC algorithm using 10 million generations and a Rtrev+G4 substitution model. Posterior probabilities for each node are indicated (values of 1 are omitted).

**Figure 5.**
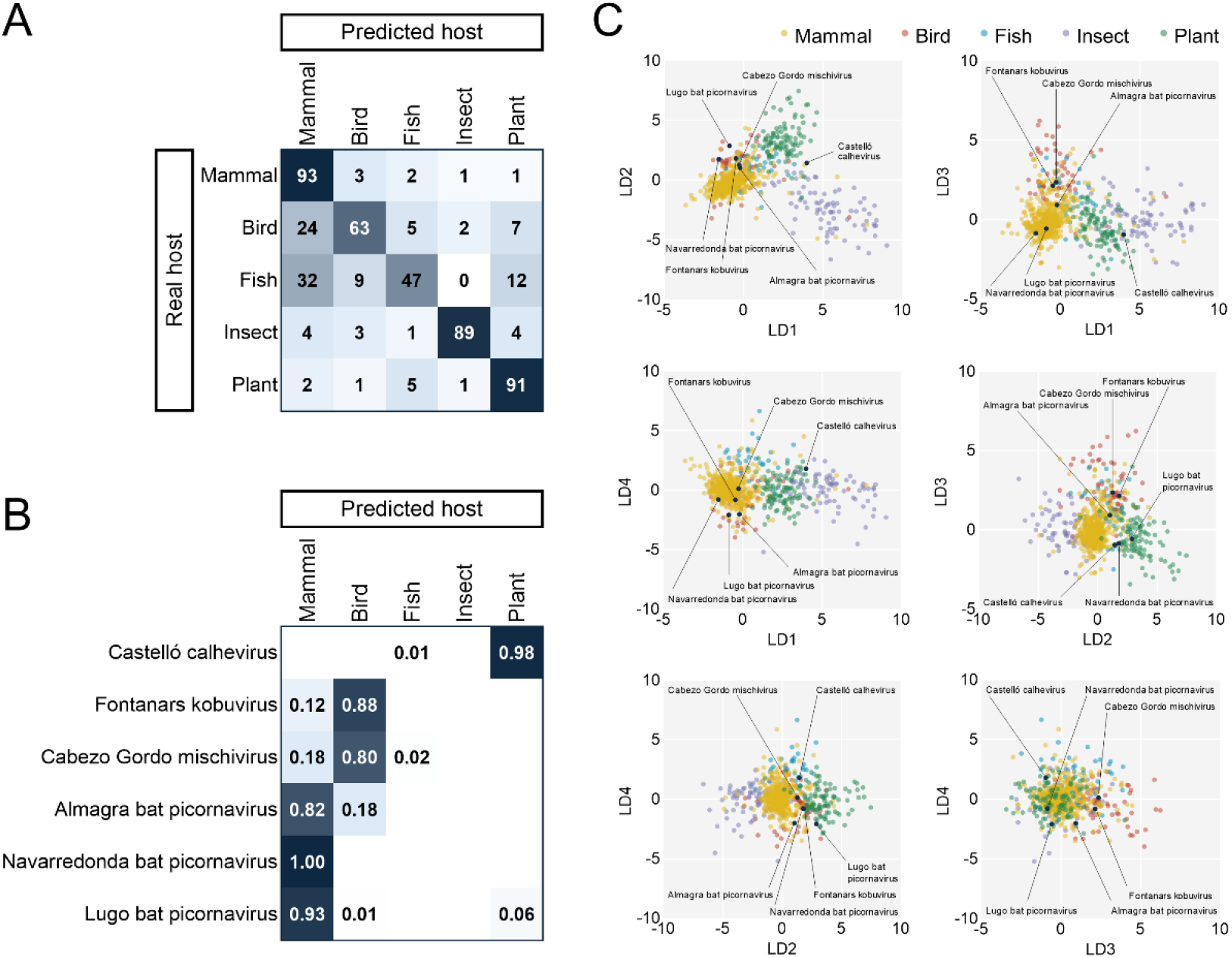
Linear discriminant analysis for the prediction of picornaviruses hosts. **A**. Confusion matrix obtained with the training dataset of 1065 viral genomes. **B**. Posterior probabilities for the classification of novel picorna-like genomes into each class (values below 0.01 are omitted). **C**. Biplots of linear discriminant analysis. Novel picorna-like sequences described in this study are shown with black dots.

**Table 2.** Picornavirus conserved sequence motifs in newly reported viruses.

| Gene | Motif | Lugo bat picornavirus | Navarredonda bat picornavirus | Almagra bat picornavirus | Cabezo Gordo bat-associated mischivirus | Fontanars bat-associated kobuvirus | Castelló calhevirus |
| --- | --- | --- | --- | --- | --- | --- | --- |
| 2A | NPGP | n.d. | n.d. | n.d. | - | n.d. | n.d. |
| 2C | GxxGxGKS | - | - | - | - | - | GxxRxGKT |
| 3C | GxCGx<sub>10-15</sub>GxH | - | GxCGx<sub>10</sub>AxHxxG | GxCGx<sub>10</sub>AxHxxG | - | - | GxSGx<sub>10</sub>GxYxxA |
| 3D | KDE | - | - | - | - | - | - |
| 3D | DxxxxD | - | - | - | - | - | - |
| 3D | PSG | n.a. | - | - | - | - | - |
| 3D | YGDD | n.a. | - | - | - | - | - |
| 3D | FLKR | n.a. | - | - | - | - | - |

n.d.: not detected; n.a.: missing due to incomplete sequence; -: identical to canonical motif

### New member of the subfamily *Ensavirinae* in *Barbastella* bats

We obtained a 7.0 kb picornavirus sequence from the feces of *Barbastella barbastellus* animals captured in Rábade and Begonte, two localities in the province of Lugo. The genome organization of this viral genome was typical of the subfamily *Ensavirinae*, a classification that was confirmed by phylogenetic analysis of the P1, 3D^pol^, and full-length genome sequences. The new sequence exhibited 63.5%, 66.4%, and 74.5% amino acid identity to Bat sapelovirus strain Henan-Mr221 (GenBank accession MF352431) in polyprotein, P1, and 2C+3CD sequences, respectively. This situates the newly described genome close to the boundary for defining a new species according to ICTV criteria for sapeloviruses (less than 70%, 64%, and 70% identity in the polyprotein, P1 and 2C+3CD amino acid sequences). While the P1 region was more closely related to that of sapelovirus and diresapivirus, the 3D^pol^ and full genome phylogenies placed it closer to parabovirus and felipivirus. We hence tentatively called this virus Lugo bat picornavirus. The cross-validated LDA model assigned a 93% probability for the virus to infect mammals, hence providing confidence that this was a true bat picornavirus and not a virus present in the diet or the environment.

### Variants of the Scotophilus kuhlii picornavirus in different *Myotis* bats

Two similar sequences of 7.7 kb sharing >94% amino acid identity in the P1 and 2C+3CD regions were obtained from a *Myotis myotis* individual captured in Navarredonda de la Rinconada (province of Salamanca) and a *M. blythii* animal dwelling in a cave called Cueva de la Almagra (Murcia) 460 km away from the other sample. Their genome structure was again typical of the subfamily *Ensavirinae*, as confirmed by phylogenetic analysis of the P1 and 3D^pol^ sequences. Phylogenetic reconstruction using full-length genomes also placed both sequences into the *Ensavirinae* subfamily. Both sequences exhibited canonical motifs in 2C and 3D proteins, but showed an alanine (GxCGx_10_AxHxxG) residue instead of the more widely conserved glycine residue in the 3C motif (GxCGx10GxHxxG). The alanine in the active site of the 3C proteinase is present in other members of the *Ensavirinae* subfamily, including all ICTV accepted species of the *Parabovirus*, *Felipivirus* and *Diresapivirus* genera. According to our phylogenetic analysis, the P1 sequences of both viruses were more closely related to those of felipivirus, boosepivirus, and parabovirus, while 3D^pol^ sequences fell into the genus *Boosepivirus*. We thus conservatively called these viruses Navarredonda bat picornavirus and Almagra bat picornavirus. The LDA model assigned a 100% probability of Navarredonda bat picornavirus to infect mammals, whereas this probability was 82% for Almagra bat coronavirus. The closest Genbank match in a blastp analysis, with >90% amino acid identity in the P1 and 2C+3CD, corresponded to Scotophilus kuhlii picornavirus, which was discovered in bat feces in 2014 in Vietnam (Genbank accession MZ544244, unpublished study). Therefore, according to ICTV guidelines, these viruses should be defined as variants of the same species. However, we notice that they were found in bats from different genera (*Scotophilus* and two *Myotis* species) and in very distant regions, suggesting a global distribution and broad host range for this virus.

### A variant of mischivirus B1 found in the feces of a *Miniopterus* bat from Murcia

A picornavirus sequence of 8.5 kb was detected in two *Miniopterus scheibersii* bats from an arid location called Cabezo Gordo, in the province of Murcia. This sequence displayed typical conserved motifs in 2C, 3C and 3D proteins, and was found to possess a NPG|P motif in the 2A polypeptide, which is present in the *Mischivirus* genus among other genera. Phylogenetic analysis of the P1 and 3D^pol^ sequences confirmed that this virus belongs to the *Mischivirus* genus, with higher posterior probabilities in the 3D^pol^ phylogeny. Phylogenetic analysis using full-length genomes also placed this sequence as a member of the *Mischivirus* genus. A blastp analysis of entire polyprotein revealed a 96.2% identity to mischivirus B1, a virus found in the feces of *M. schreibersii* in Hungary [47], indicating that our virus constitutes a variant of mischivirus B1. Genome composition analysis identified birds as the most likely host of the newly described virus with an 80% probability, versus only a 18% for mammals. Thus, an environmental origin from bird material cannot be ruled out, although it should be noted that the LDA model showed poor ability to discern between different types of vertebrate hosts. We hence called this virus Cabezo Gordo bat-associated mischivirus. In any case, it is highly unlikely that this virus came from the diet since *M. scheibersii* is an insectivorous bat.

### A divergent variant of Wenzhou bat picornavirus 7 found in *Pipistrellus* bats from Valencia

Another picornavirus sequence of 8.5 kb was found in the feces of four *Pipistrellus kuhlii* individuals from different pools obtained in Fontanars, in the province of Valencia. The sequence exhibited a genome layout matching the *Kobuvirus* genus, and protein 2A displayed an H-box (HWAV) upstream from a NC-box that is also typical of this genus [48]. Phylogenetic analysis with P1, 3D^pol^, and full-length genome sequences confirmed that this was a member of the *Kobuvirus* genus. Genome composition analysis identified birds as the most likely hosts with an 88% probability, versus only a 12% for mammals. Thus, we tentatively called this virus Fontanars bat-associated kobuvirus. As above, it is unlikely that this virus came from the diet, since *P. kuhlii* is also an insectivorous bat. An environmental origin from bird material cannot be discarded but appears to be less likely than an incorrect classification given the poor ability of the LDA model to discern between avian and mammalian hosts. The closest ICTV-accepted species in the phylogeny was aichivirus F1, but with a high divergence. The best blastp hit corresponded to Wenzhou bat picornavirus 7 (Genbank accession OQ716048), a picornavirus recently discovered in *P. abramus* from China [49]. These two sequences shared 72.8% sequence identity across the whole polyprotein, while percentage identities for P1 and 2C+3CD were 77.6% and 74.7%, respectively. Hence, according to ICTV criteria, our sequence and Wenzhou bat picornavirus 7 should be considered members of the same species. However, they are relatively divergent and were found in different *Pipistrellus* species from very distant regions, suggesting that this virus is present globally, displays high genetic diversity, and shows interspecies transmissibility.

### A plant-like calhevirus found in a *Tadarida* bat from Castelló

A 10 kb picorna-like sequence was obtained from the feces of *Tadarida teniotis* bats captured in Castelló. This sequence exhibited quite dissimilar motifs in the 2C and 3C proteins compared to picornaviruses, showing a GxxRxGKT motif in the 2C protein instead of the widely conserved GxxGxGKS, and a serine residue in the catalytic site of the 3C protease (GxSGx_10_GxYxxA) instead of the typical cysteine. The best hit in a blastn analysis using the entire sequence (68.7% nucleotide identity along a 28% coverage span) corresponded to Jinngmen bat picorna-like virus 1, a virus recently discovered in *Myotis chinensis* [49]. All other top hits belonged to calhevirus sequences [26,50], with similar nucleotide identities and slightly lower coverages. The non-structural ORF1 of our sequence shared a 51.0% amino acid identity to that of Jingmen bat picorna-like virus 1, whereas the capsid polyproteins exhibited 53.6%, indicating that this should be a novel calhevirus species, which we called Castelló calhevirus. Genome composition analysis identified plants as the most likely hosts with 98% probability. *T. teniotis* is yet another insectivorous bat, but ingestion of plant material associated with insects cannot be discarded.

## Conclusions

Investigation of the virome of 23 different bat species across Spain’s geography led to the identification of six complete or quasi-complete new viral genomes belonging to the *Picornavirales* order. Our results provide information about the host, geographical range, and genetic diversity of bat-associated picornaviruses. Two of the described viral genomes may constitute new species according to ICTV criteria. One of these viruses, Lugo bat picornavirus, is likely to infect bats, whereas the other is a divergent calhevirus that may infect plants, according to genome composition analysis. Previously annotated calheviruses were isolated from human, shrew, and rat feces, and were inferred to infect arthropod hosts using genome composition analysis [26,50,51]. The newly described Castelló calhevirus might constitute the first member of this family infecting plants.

We also found a new kobuvirus in multiple *Pipistrellus kuhlii* individuals, suggesting that this was a prevalent virus amongst the local bat population of Valencia, although this abundance might be related to a certain outbreak in these bats. Importantly, members of the *Kobuvirus* genus can cause gastroenteritis and diarrhea in humans and other mammals such as pigs [52,53]. The newly detected kobovirus was highly divergent from its closest ICTV-accepted relative (Aichivirus F2), and was closer to a virus recently discovered in a different species of the same bat genus in China, the similarity between these two viruses being close to the species demarcation threshold (ca. 70% amino acid identity).

Three additional viral genomes belonged more clearly to previously described species. Two of these genomes, obtained from different species of *Myotis* in different regions of Spain, showed >90% amino acid identity with a picornavirus described in 2014 in lesser Asiatic yellow bats (*Scotophilus kuhlii*) from Vietnam, showing that these sequences pertain to a geographically widespread and genetically diverse picornavirus capable of infecting bats from different genera. Another picornavirus genome found in the feces of *M. schreibersii* was shown to be a variant of mischivirus B1 with 96.2% amino acid identity to another viral sequence obtained from the same bat species in Hungary in 2013 [47]. Mischiviruses have also been reported in canine and shrew-derived samples, suggesting that this genus can infect a broad range of mammals.

Both the Fontanars bat-associated kobuvirus and the Cabezo Gordo bat-associated mischivirus most probable hosts were predicted to be birds, rather than mammals, despite being isolated from bat fecal samples. Although dietary or environmental origins are unlikely considering their insectivorous nature, such origins cannot be fully disregarded. For this purpose, further studies into the virome of both Spanish bat and bird communities could assist in unraveling the tropism range of these newly identified viruses. In that direction, further research effort is warranted to confirm the predicted bird tropism, which could increase their zoonotic potential, also considering the expanded cross-species infectivity and transmissibility between bats species reported here for the Fontanars-bat associated kobuvirus, and Navarredonda and Almagra bat picornaviruses.

Overall, our results show that picornaviruses are distributed widely in the Spanish bat population. For some of these picornaviruses, we found evidence that variants of the same virus can be found in distant parts of the world and can infect divergent bat species and, potentially, other vertebrates. In light of these findings, special protection of the natural habitats of Spanish bats is warranted to minimize zoonosis risk. Further research efforts should be implemented towards characterizing the abundance of picornaviruses and other viral families in different types of hosts and different regions of the world in order to obtain a more complete picture of their diversity, cross-species transmissibility, and associated human spillover risks.

## Declarations

### Ethics approval and consent to participate

Samples consisted of feces from wild animals captured using nylon mist nets or a harp trap. Bats were kept briefly in cotton bags until fresh fecal samples were obtained. According to the European Directive regulating the protection of animals used for scientific purposes (2010/62/EU, Article 1), subsequently transposed into Spanish legislation (Royal Decree 53/213, 1 February, Article 2), procedures used in this study are not subject to the condition of animal experimentation and therefore do not require a IACUC approval document, but specifically a permit from the competent regional authority. The necessary permits from the Generalitat Valenciana for the sampling of wild bats were granted under Exp. 2022-VS (FAU22_009).

### Consent for publication

not applicable.

### Availability of data and materials

raw sequence reads were deposited in the GenBank Sequence Read Archive under accession number SRR27912329, SRR27912332, SRR27912333, SRR27912339, SRR27912350, SRR27912351, and SRR27962820-SRR27962839. The viral contigs corresponding to complete or nearly complete genomes were deposited in Genbank under the accession numbers provided in Table 1.

## Authors’ contributions

MCS integrated data analyses and drafted the manuscript. JB obtained viral contigs and analyzed data. AV obtained bat feces samples. IAM performed LDA. RMR prepared samples for sequencing. CST contributed to viral detection by PCR. JSM supervised work and obtained bat feces samples. JMC obtained funding, supervised work, and revised the manuscript. RS obtained funding, supervised work, and co-wrote the manuscript.

## Acknowledgements

we thank members of the Virus Evolution lab for helpful comments and suggestions.

## Supplementary material

**Table S1.** Detailed list of specimens and samples examined in this study. Detailed list of sampled specimens, including collection date, species name, age, sex, type of trap used and pooling.

**Table S2.** Illumina reads and viral contigs obtained. Sequencing data for each pool, including raw and clean reads, number of contigs, number of viral contigs, and complete or high-quality RNA genomes.

**Table S3.** Specific primers used for PCR confirmation. Target viral genome and target genomic region, 5’ 3’ sequence, and amplicon length are indicated.

**Table S4.** Nucleotide composition-based linear discriminant model. Detailed information for the model used, including accession numbers, reported host, mono- and dinucleotide frequencies, dinucleotide bias, and resulting linear discriminants.

**Figure S1.** Phylogenetic tree of the Picornaviridae family for the complete polyprotein. The newly described viruses are placed in the context of ICTV-approved viruses. Ampivirus A1 (accession KP770140) was used as outgroup. Phylogenetic inference was done using maximum likelihood and a GTR+I+G4+F substitution model. Bootstrap values are shown in grey for each node.

