## Supplementary figures and images for "Genetic diversity and cross-species transmissibility of bat-associated picornaviruses from Spain"

### Supplemental Figure 1

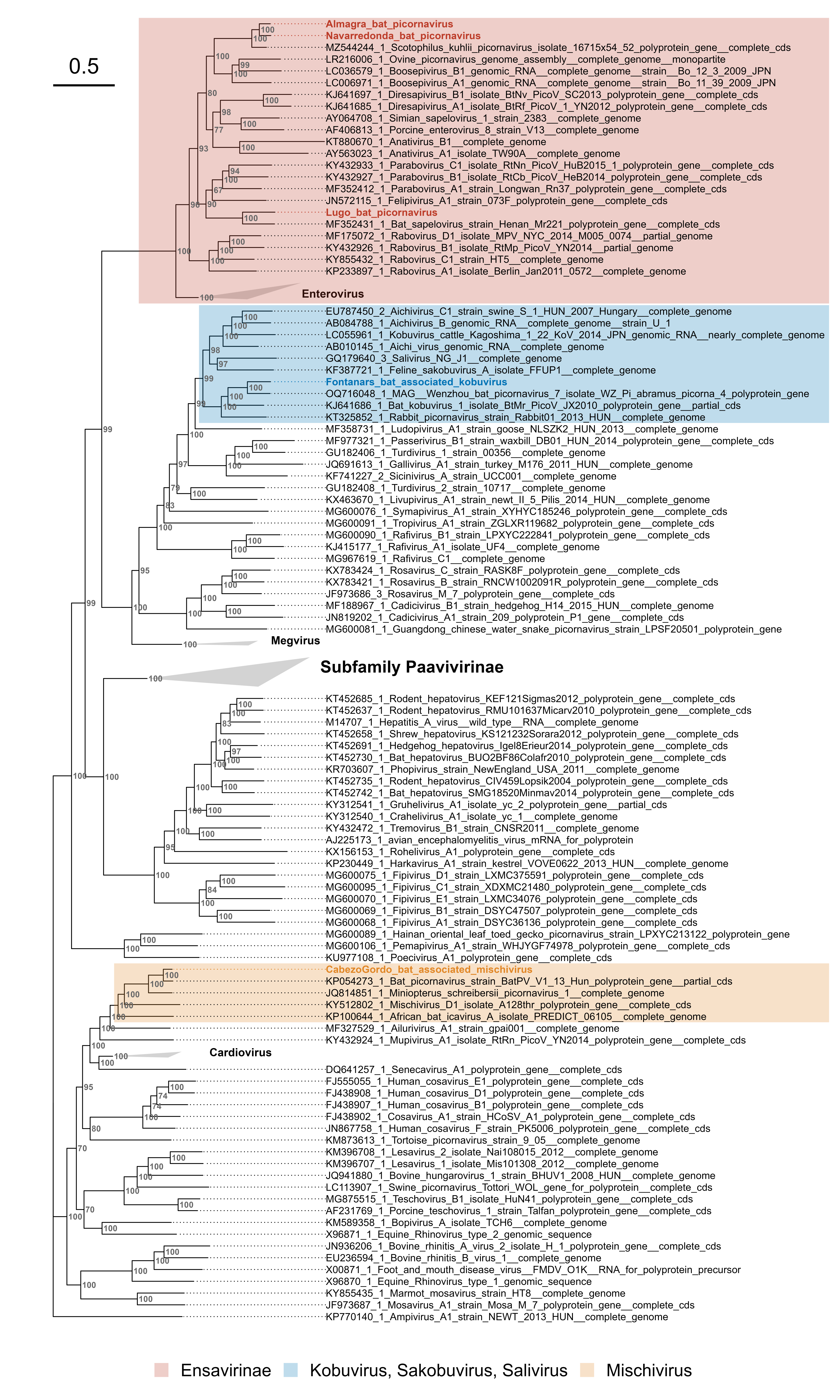
